# MR-ARFI and Beyond: Characterizing the Temporal Dynamics of Acoustic Radiation Force in Human Brain

**DOI:** 10.64898/2026.09.29.755540

**Authors:** Morteza Mohammadjavadi, Kim Butts Pauly, Gary H. Glover

## Abstract

**Purpose:** Magnetic Resonance Acoustic Radiation Force Imaging (MR-ARFI) encodes focused ultrasound (FUS)-induced tissue displacements to MR phase shifts. Because these phase shifts are small, optimizing displacement sensitivity and minimizing motion artifacts requires characterization of the temporal dynamics of acoustic radiation force (ARF). Beyond sequence design, such characterization enables MR-ARFI to serve as a probe of brain viscoelastic behavior and can inform the design of therapeutic ultrasound pulsing schemes by quantifying how rapidly tissue deforms and relaxes.

**Methods:** We developed a dynamic model assuming a gamma-variate impulse response for ARF(t), with time constant (τ) to be estimated. A single-shot spin echo ARFI acquisition with a bipolar motion encoding gradient (MEG) was implemented, incorporating retrospective corrections for head motion and physiological noise. A 500 kHz FUS pulse was applied in on/off blocks while systematically varying the timing between the MEG and sonication. The resulting phase time series were fit to the model to estimate τ in five human participants. To assess sensitivity to tissue stiffness, τ was additionally measured in two customized gelatin phantoms of differing stiffness, characterized independently by MR elastography (MRE).

**Results:** The average measured time constant in human participants was τ= 3.7 ± 0.27 ms. Comparing the two phantoms, the stiffer phantom exhibited a shorter τ (3.0–3.5 ms) than the softer phantom (4.0–4.5 ms), which tracked the MRE-derived Kelvin-Voigt viscoelastic time constants. In addition, the peak ARFI displacement was lower in the stiffer phantom (0.17 vs 0.74 μm for the less stiff phantom), and a broader displacement focal spot size (FWHM 22 mm vs 15 mm).

**Conclusion:** This time-series approach with motion and physiological noise corrections validates an MR-based technique capable of characterizing the brain’s temporal response to sonication-induced mechanical forces, providing a foundation for optimizing the design of ultrasound pulsing schemes for ARFI, as well as for future studies of brain biomechanics.

## Introduction

Magnetic Resonance Acoustic Radiation Force Imaging (MR-ARFI) provides a noninvasive method to visualize focused ultrasound (FUS) fields by encoding radiation force–induced tissue displacements into MR phase using motion encoding gradients (MEGs)[1], [2], [3], [4]. MR-ARFI is increasingly used for focal verification in transcranial ultrasound applications, where accurate localization is essential for both therapeutic and neuromodulatory studies. Its performance depends on achieving high displacement sensitivity while maintaining low ultrasound exposure and robust acquisition efficiency. In prior work[5], we developed a single-shot spiral MR-ARFI sequence that enabled in vivo ARFI mapping in the human brain with substantially improved signal-to-noise ratio and reduced ultrasound exposure compared to conventional multi-shot acquisitions[3], [6], [7]. This approach demonstrated rapid, motion-robust ARFI suitable for clinical applications such as targeting of FUS for neuromodulation[8] or ablative therapy[9].

While MR-ARFI is well established for localizing the focus, the motion it encodes is not merely linear displacement. Tissue deforms during the applied ultrasound force and relaxes afterward, following a characteristic time course determined by the tissue’s mechanical properties. This temporal response carries information beyond focal position. Prior work characterized the ARF-induced displacement time constant in post-mortem pig brain using unipolar MEGs and an exponential model[10], but the temporal dynamics of ARF-induced motion have not been characterized in the living human brain[11].

Characterizing this temporal response serves three purposes. First, ARFI sequence optimization and dose reduction is enabled: precise knowledge of the displacement onset and peak facilitates optimized MEG–FUS timing, improving phase sensitivity and minimizing acoustic exposure. Second, noninvasive probing of tissue biomechanics can be explored. Because the temporal profile of ARF-induced deformation in brain tissue reflects viscoelastic behavior, it positions MR-ARFI as a tool for mechanical characterization in addition to focal localization. Brain tissue is mechanically heterogeneous, with shear stiffness varying across regions as demonstrated by MR elastography (MRE)[12], [13], [14], [15]. Therefore, a measure of the mechanical response obtained directly from the localization sequence would be valuable. A third application where a knowledge of the ARF temporal dynamics is important is informing therapeutic sonication design. Because the displacement response depends on how rapidly tissue deforms and relaxes, knowledge of the time constant can guide the choice of sonication pulse duration and pulse repetition frequency (PRF) to achieve a desired mechanical effect.

In the present study, we extend our single-shot spiral ARFI framework[5] to characterize the temporal response of brain tissue to acoustic radiation force. We apply a single bipolar MEG and a brief FUS pulse while systematically varying their relative timing, and model the resulting displacement using a gamma-variate impulse response with a time constant τ [16] [17] that characterizes the temporal dynamics of ARF. We measure τ in the human brain, assess its sensitivity to tissue stiffness using gelatin phantoms characterized independently by MRE, and illustrate through simulation how it informs ultrasound pulse design. The high degree to which the measured ARFI response aligns with that predicted by the gamma variate model is taken as a measure of model validity. By introducing displacement modeling into MR-ARFI, this work advances the technique from static displacement mapping towards dynamic assessment of tissue mechanics in the human brain.

## Methods and Materials

### Theory: Model of ARF

The time-dependent tissue displacement Δ*z*(*t*) during sonication is modeled as a convolution of the sonication pulse waveform Fus(t) with an acoustic radiation force impulse response function Arf(t) as

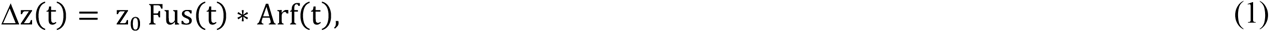

where *z*_0_ is a constant and *Fus(t)* represents the acoustic radiation force envelope; in this experiments it is a rectangular pulse of duration *T*_*fus*_ (constant-amplitude sonication), though the convolution model (Eq. 1) applies to any known force waveform. Motivated by the shape of the measured phase response and consistent with prior tissue displacement studies[16], [17], we model the impulse response function Arf(t) as a first-order gamma-variate function:

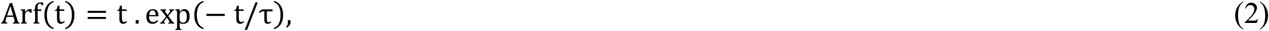

where τ is the characteristic time constant that is to be determined. To measure τ, we employ a pulse sequence with a bipolar MEG[18] and, in separate scans, shift the start of the FUS relative to the MEGs by an amount *T*_*off*_ as shown in Fig. 1. The resulting angular frequency shift of spins in the sonicated region is

**Figure 1.**
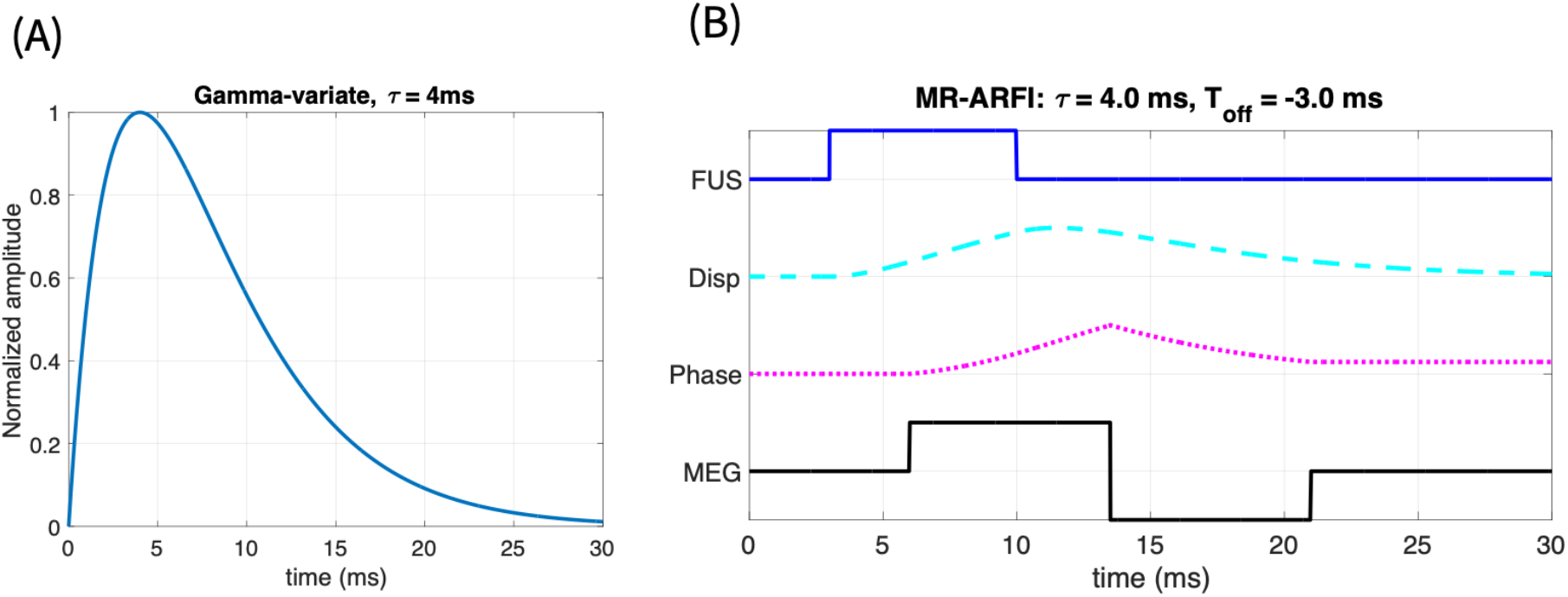
Model for characterizing the impulse response function Arf(t)’s time constant τ. (A) Assumed gamma variate impulse response function, with τ= 4ms. (B) Bipolar MEGs (black), FUS pulse (blue) starting at time *T*_*off*_ relative to the start of MEGs, calculated displacement (cyan) from Eq. 1, and MR phase shift (magenta), Eq. 4. As *T*_*off*_ is changed between separate measurements, the ending accumulate phase Δϕ_meas_ is fit to the model.

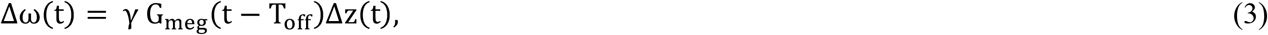

where *G*_*meg*_(t) is the MEG waveform and γ is the gyromagnetic ratio for protons. The phase Δϕ accumulated during the MEG duration T_meg_ is

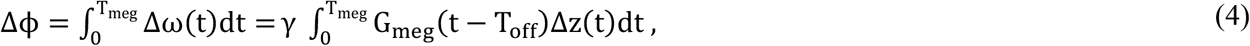

where t = 0 is the start of the MEGs. From Eq. 1, the tissue displacement is a function of time as well as details of the MEGs and *T*_*off*_. Figure 1 demonstrates the model for an assumed value of τ = 4 ms. Note that tissue displacement can arise before the MEGs start and can continue beyond their end, but the phase accumulation Δϕ*meas* only occurs during the MEGs. Evaluation of Eqs. 1 and 4 for a range of *T*_*off*_ and τ forms the basis for estimating τ from measurements of the total phase shift, Δϕ*meas*(*T*_*off*_).

### MR data acquisition

Scans were acquired on either of two 3T scanners labeled MR1 (GE Discovery 750, Milwaukee Wi., 8 channel head coil, 50 mT/m MEG gradients) and MR2 (GE Premier, Milwaukee Wi., 48 channel head coil, 70 mT/m MEG gradients) as shown in Table 1. Aside from differing MEG amplitudes, the pulse sequence design and other parameters were identical. Human participants 1–2 were scanned on MR1, while the custom-built phantoms and human subjects 3–5 were scanned on MR2. The pulse sequence is shown in Fig. 2. The images were acquired with in-plane resolution of 2.75×2.75 mm^2^ in 3 slices with thickness 4mm, and were reconstructed into 1.72×1.72×4 mm voxels, echo time TE = 60 ms, TR = 800ms, and either T_meg_/T_fus_ = (10/6 ms) or (15/7 ms).

**Table 1.** Human ARFI scan parameters and results, including the correlation between measured phase as a function of T_off_ with modeled phase for the best fit τ value.

| Participant | $T_{meg}$, ms | $T_{Fus}$, ms | $\tau$, ms | $R^2$ (Correlation of Model vs. Measured phase), $Df(T_{off})$ | $p$ | MR/<br>design |
| --- | --- | --- | --- | --- | --- | --- |
| 1 | 10 | 6 | 4.0 | 0.98 | < 0.0001 | MR1/D1 |
| 2 | 15 | 7 | 4.0 | 0.93 | < 0.0021 | MR1/D1 |
| 3 | 15 | 7 | 3.5 | 0.96 | <0.0001 | MR2/D2 |
| 4 | 15 | 7 | 3.5 | 0.99 | <0.0001 | MR2/D2 |
| 5 | 15 | 7 | 3.5 | 0.99 | <0.0001 | MR2/D2 |
| Average | | | $3.70 \pm 0.27$ | $0.97 \pm 0.02$ | | |

**Figure 2.**
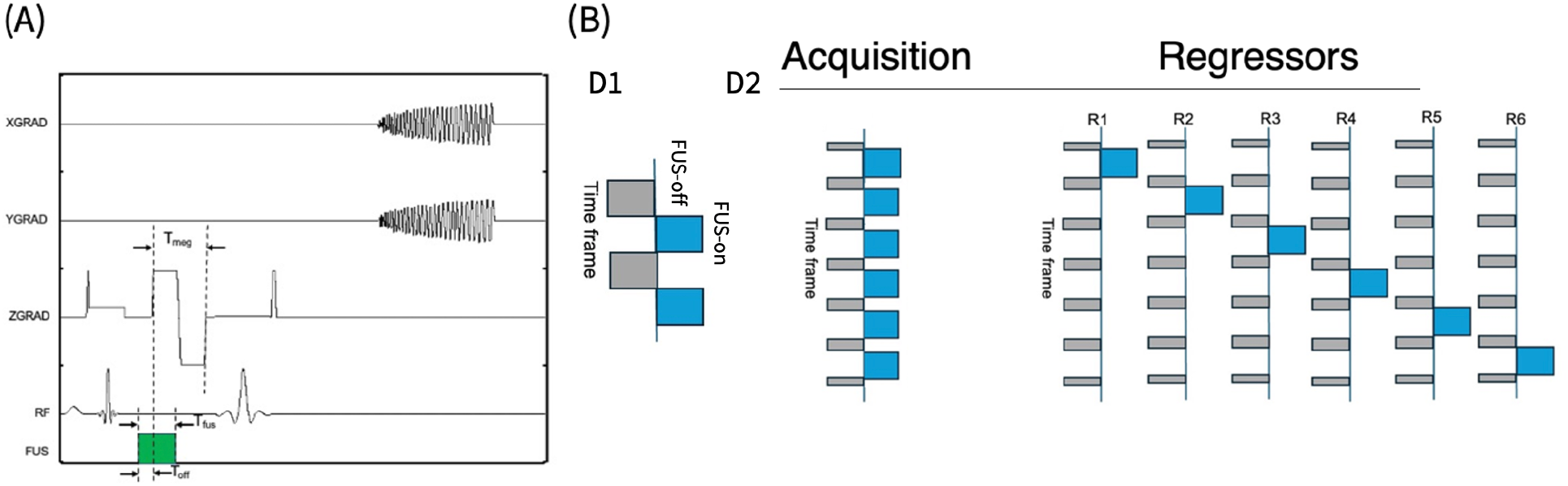
Spin-echo pulse sequence using single-shot spiral readout to estimate Arf(t) τ as *T*_*off*_ is varied. Spiral trajectories are more efficient than EPI, resulting in shorter readout duration (22ms vs. 34ms) to limit the deleterious impacts of motion. The initial RF pulse and subsequent Z-gradient before the 90° pulse are included for fat saturation (A) Design matrices for GLM analysis. Grey boxes indicate baseline (no-FUS) blocks of time frames with weight - 1; blue box indicates FUS-on blocks with weight +1. No-box time frames have weight 0. (D1) Design D1 has one regressor, and 25 time frames in each block for a scan. Scan is repeated 6X with different values of T_off_ to obtain the full measurement set. (D2) Design D2 has 6 regressors (R1-R6) for the single scan, with different values of T_off_ employed for each of the 6 sections of the scan. FUS-on blocks are 48 time frames long; inner off-blocks are 8 time frames long; first and last off-blocks total 8 time frames for a total of 336 time frames in the scan. This single-scan design reuses the 48 total baseline time frames and is thereby more efficient (in time and in total ultrasound dose) than D1 repeated for a total of 600 time frames.

Two timeseries designs for the application of FUS pulses were employed (Fig 2). In the first (D1, used with participants 1 & 2), 100 time frames (TRs) were acquired in FUS “off-on-off-on” blocks of 25 TRs/block, in each of 6 scans with differing *T*_*off*_ values. This design required a total of 600 TRs. The second design (D2, used with participants 3 - 5) is performed in one scan having 6 “on” blocks of 48 TRs each with the different *T*_*off*_ values, interspersed before and after with “off” blocks shared in the GLM for all 6 “on” conditions. This design has 48 “on” and 48 “off” TRs for the 6 *T*_*off*_ conditions for a total of 336 TRs, which substantially reduces the total scan time (and FUS dose) while nearly maintaining the same number of degrees of freedom in the model (Figure 2).

Utilizing a bipolar MEG creates a wide dynamic range of phase shift as *T*_*off*_ is varied. Second-order shimming was employed before ARFI scans to diminish off-resonance generated by the FUS transducer and participant’s head. Anatomic imaging used T2-weighted fast spin-echo (TR/TE = 3000/68 ms).

### Sonication

Different 500 kHz focused ultrasound systems were employed with each scanner. In MR1, a 4 element annular array (Neurofus System, UK) was used with a free-water peak pressure of approximately 1.48 MPa

(*I*_*sppa*_ = 68 *W*/*cm*^2^ at 60 mm natural focal depth), as measured with a calibrated hydrophone (Onda, Sunnyvale, USA). In MR2, a 64 element Image-Guided-Therapy system (Image Guided Therapy, Paris, France) was used with calibrated free-water peak pressure of approximately 1.34 MPa at 65 mm focal depth. In all cases the ultrasound pulse was applied during half the time frames with the sequence in Fig. 1 for each of the three slices.

### Human scans

Five human subjects were scanned after obtaining informed consent as approved by the Stanford IRB. The transducer was coupled to the head using ultrasound gel (Aquasonic 100 Ultrasonic gel, Parker Laboratories, NJ, USA) and a 10 mm thick gel-pad (Aquaflex, Parker Laboratories, NJ, USA). The depth of sonication focus was 60 or 65 mm, which reached the left thalamus in all participants. No attempt was made to precisely target a specific brain region in this study. The first participant was scanned with shorter MEGs and FUS (T_meg_/T_fus_ = 10/6 ms), while participants 2-5 had scans utilizing T_meg_/T_fus_ = 15/7 ms which was more optimum for the longer observed human average τ = 3.7ms.

### Gelatin Phantom Fabrication

Custom phantoms were fabricated using a gelatin–evaporated milk formulation. Two homogeneous phantoms were prepared using 175-bloom porcine gelatin (Sigma-Aldrich, St. Louis, MO) with different gelatin concentrations to produce distinct mechanical stiffnesses while maintaining the same liquid composition. Each phantom consisted of 350 mL deionized water and 350 mL evaporated milk (Carnation, Nestlé), with gelatin concentrations of 25 g and 50 g (Phantom 25 and Phantom 50, respectively). A silicone defoamer (Antifoam A, Sigma-Aldrich) was added to minimize air bubbles during fabrication. Gelatin was dispersed in deionized water, dissolved by heating in a water bath, followed by incorporation of the evaporated milk. The mixtures were poured into molds, refrigerated overnight to solidify, and allowed to equilibrate to room temperature before imaging and acoustic measurements [19], [20].

### Gelatin Phantom Characterization

To assess the sensitivity of the ARFI τ to tissue stiffness, the viscoelastic properties of the two gelatin phantoms were characterized independently by MR elastography (MRE). MRE was performed on a 3T GE Signa Premier scanner (GE Healthcare, Milwaukee, WI) using a 48-channel head coil. Shear waves were induced in the phantom by a pneumatic passive driver (Mayo Clinic, Rochester, MN). Imaging parameters were: acquisition matrix = 128 × 128, field of view = 22 × 22 cm, slice thickness = 3 mm. Storage (G′) and loss (G″) moduli were measured at five actuation frequencies (40, 60, 80, 100, and 120 Hz); the repetition and echo times were automatically adjusted at each frequency to synchronize the motion-encoding gradients to the vibration period (TR = 2059–2275 ms, TE = 61–80 ms).

For each phantom, the phantom region was identified from the MRE magnitude image by thresholding at 25% of the maximum intensity and retaining the largest connected component; a circular Region of Interest (ROI) of 10 mm radius, centered on the phantom centroid, was then intersected with this mask. G′ and G″ were averaged within this ROI across six slices.

To match the definition used for the reference brain data (Hiscox et al,), the complex shear modulus magnitude was computed at each frequency, and reformulated to shear stiffness[15] μ as

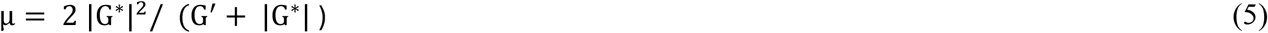

where the complex shear modulus is G^*^. To enable comparison with the human brain literature value, the phantom shear stiffness was interpolated to 50 Hz, matching the frequency of the reference brain MRE data[15]. The static (peak) ARFI displacement in each phantom was also obtained from the encoded phase at the focus. To relate the frequency-dependent MRE moduli to a viscoelastic time constant comparable to the ARFI measurement, we fit the measured moduli to a Kelvin-Voigt (KV) model[13], [21], in which a spring (stiffness μ) and dashpot (viscosity η) act in parallel, giving a complex shear modulus G^*^ = μ + iωη, where ω is the angular frequency. Under this model the storage modulus equals the elastic stiffness (G′ = μ) and the loss modulus is proportional to frequency (G″ = ωη), so the viscoelastic time constant is:

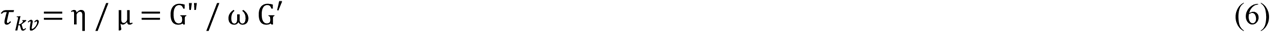

The stiffness μ was taken as the mean storage modulus across frequencies, and the viscosity η was obtained from the slope of G″ versus angular frequency by least-squares regression. This yields a single, frequency-independent time constant τ_kv_for each phantom that reflects the ratio of viscous to elastic behavior, providing a direct viscoelastic analogue to the ARFI-derived time constant. The peak ARFI displacement in each phantom was obtained from the encoded phase at the focus, converting phase to displacement using the known MEG amplitude, duration, and gyromagnetic ratio. The focal spot size was quantified as the full width at half maximum (FWHM) of the displacement profile through the focal peak, using both a direct profile measurement and a two-dimensional Gaussian fit.

### Data analysis

The timeseries of complex images is processed using a general linear model (GLM) to generate statistical maps of the difference between phase with FUS on and off states, as shown in Fig. 2. The GLM for each value of *T*_*off*_ is

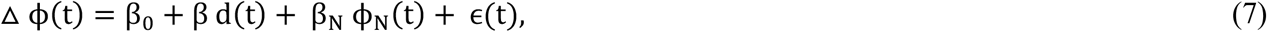

where *d(t)* is the off-on block design, D1 or D2 (-1 for off, +1 for on), β is the amplitude of the design contained in the phase data △ ϕ(*t*), β_0_ is the average phase, β_*N*_ is the amplitude of nuisance regressor for bulk brain phase ϕ_*N*_(*t*), and *e(t)* is residual error not explained by the other regressors. β is used to generate T-scores of sonication and to estimate the Arf(t) dynamics. The ARFI maps were generated using the same method in Mohammadjavadi et al, 2025[5].

For each scan with a given T_off_, the ARFI signal was averaged over a focal region of interest (ROI). The ROI was first defined as the connected cluster of voxels significant at p < 0.03 (t ≥ 1.9, 56 degrees of freedom, one-sided) containing the focal peak (located as the maximum of the smoothed t-map within the brain interior). For the phantoms, because a stiffer phantom can produce a larger significant region while the physical ultrasound focus is unchanged, the ROI was additionally limited to the 27 highest-t voxels, corresponding to the 10 mm focal-spot size at the 1.72 mm in-plane resolution, so that the ROI reflected the physical beam dimension and was comparable across phantom measurements. This size limit was not applied to the human data, where the low-SNR ARFI response was already smaller than the focal-spot size and thus fully contained within the significance-defined cluster.

Within this ROI, the β values were averaged for each T_off_ to yield the measured phase-versus-offset curve. To estimate time constant τ, the measured curve was compared with model predictions computed from Eqs. 1–4 over a range of candidate time constants (τ = 1.0–6.0 ms in 0.5 ms increments). For each candidate τ, a single amplitude scale factor was determined by least squares, and the τ giving the smallest residual between the measured and modeled curves was selected as the best fit. For the phantoms, the fitted τ was robust to the significance threshold and ROI size, varying by at most one grid increment (0.5 ms) across a range of thresholds.

### Pulse Design Simulation

Because the displacement response is governed by the time constant τ (Eqs. 1–2), knowledge of τ informs the choice of sonication pulse parameters. For a single rectangular FUS pulse of duration T_fus_ convolved with the gamma-variate impulse response, the peak displacement Z_peak_, as a fraction of its saturation value Z_max_ (the maximum peak displacement achievable in the long-pulse limit, T ≫ τ), is

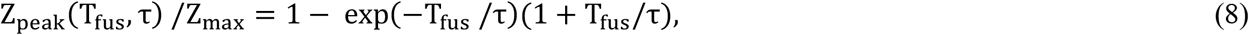

which depends only on the ratio *T*_*fus*_/τ. Two aspects of pulse design were examined by simulation using the gamma-variate model: (i) the peak displacement of a single rectangular FUS pulse as a function of *T*_*fus*_/τ and (ii) for pulse trains at fixed duty cycle, the displacement time course for several pulse repetition frequencies (PRFs) using the brain τ (τ = 4 ms). These simulations used the same convolution model (Eq. 1) employed for τ estimation.

## Results

### Human

For the first two participants, *β* maps for all *T*_*off*_ values, along with the extracted β values overlaid on the model are shown in Fig. 3. Substantial differences are observed between the models and corresponding measurements in panels (A) versus (B), reflecting the different *T*_*off*_ and *T*_*fus*_ parameters used, as summarized in Table 1. Despite these variations in timing, the measured curves for participants 1 and 5 were best described by the models with nearly identical τ, 4.0 and 3.5 ms, respectively. Results for the other participants were very similar to those in Fig. 3 and are therefore not shown. Table 1 summarizes the parameters and results for all participants, with the last column describing which of the two scanners and which design (Fig. 3) were employed. The average τ = 3.7 ± 0.27 ms. The table includes *R*^*2*^ *and p values* for the correlation between measured and modeled phase, indicating very high agreement in all cases despite changes in the parameters, scanner and experiment design.

**Figure 3.**
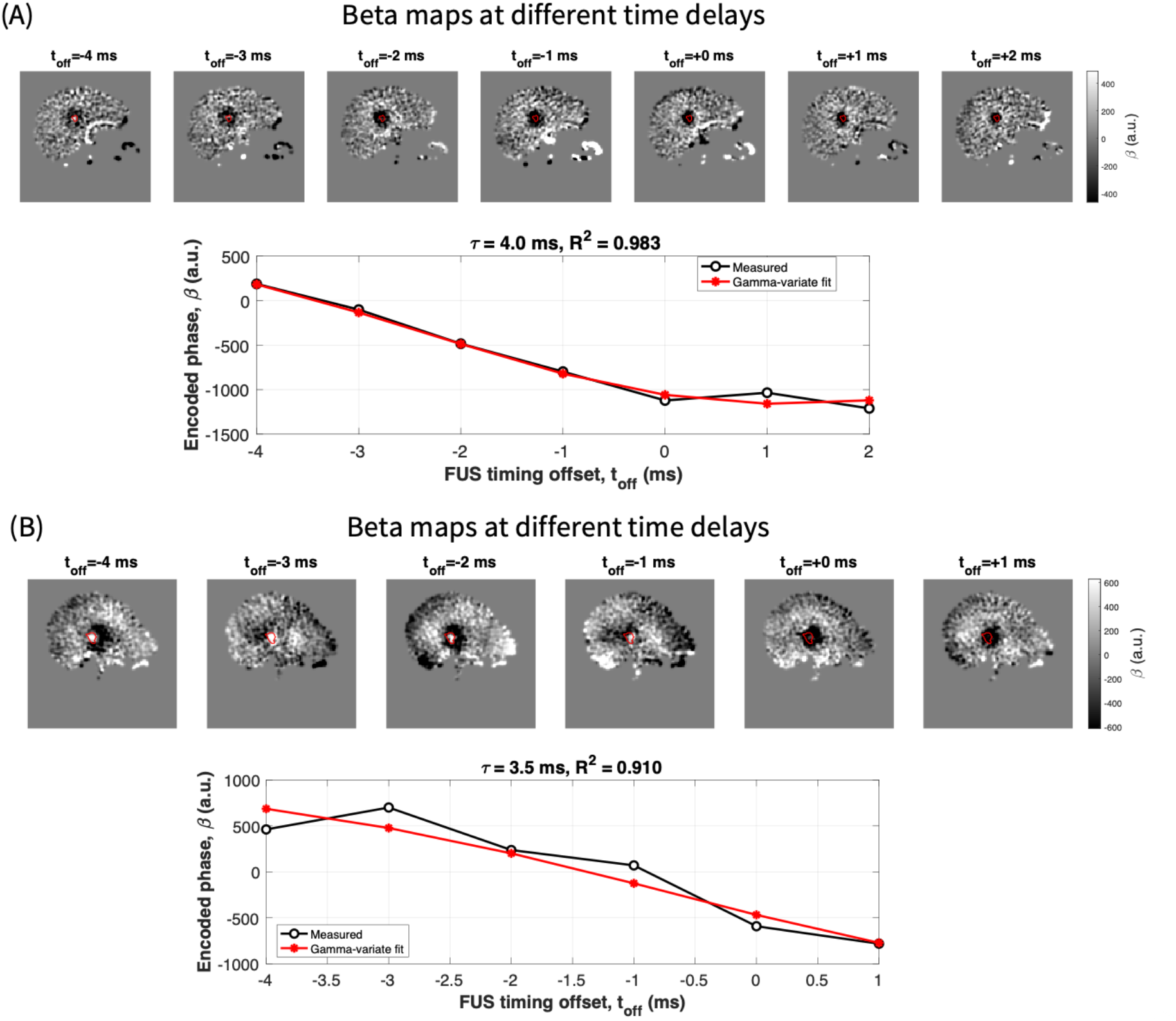
Representative scan and model results for two human participants. β maps are shown for the slice at the center of the FUS focus. Modeled phase versus T_off_ is plotted for 4 values of τ (solid multi-colored curves) with the measured β values overlaid. Despite differences in acquisition parameters T_meg_ and T_fus_ and experiment design, the measured curves for both participants are well fit (R^2^ >0.9) by the models with nearly identical τ.

### Sensitivity to Tissue Stiffness

Phantom 50 exhibited higher storage and loss moduli than Phantom 25 across all measured frequencies, confirming it as the stiffer phantom (Fig. 4). Interpolated to 50 Hz frequency, the measured shear stiffness was 1.78 kPa and 2.97 kPa for Phantom 25 and Phantom 50, respectively.

**Figure 4.**
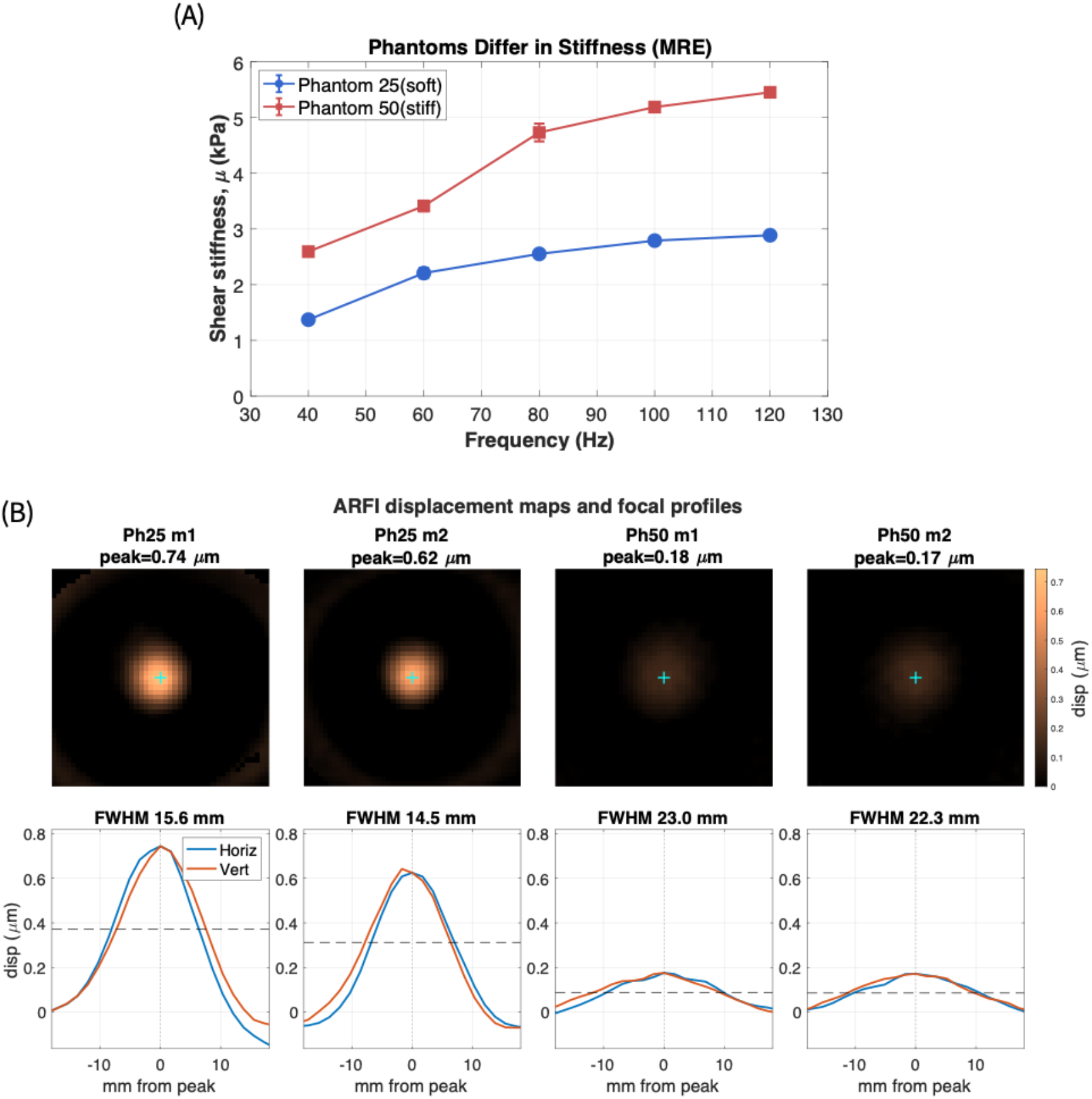
MRE shear stiffness versus frequency for the two custom gelatin phantoms. Phantom 50 (stiff) exceeds Phantom 25 (soft) at all frequencies; error bars are the standard error across six slices (panel A). ARFI displacement maps (top) and focal profiles (bottom) for two measurements each of the soft (Phantom 25) and stiff (Phantom 50) phantoms. The stiffer phantom exhibits markedly lower peak displacement (0.17–0.18 vs 0.62–0.74 μm) and a broader focal displacement profile (FWHM 22–23 vs 14–16 mm) (panel B).

The two phantoms also differed markedly in their ARFI displacement (Fig. 4). Across two measurements each, the softer Phantom 25 exhibited a peak displacement of 0.62–0.74 μm, while the stiffer Phantom 50 showed a substantially lower peak displacement of 0.17–0.18 μm, consistent with reduced deformation in the stiffer medium under the same sonication. In addition, the stiffer phantom exhibited a broader displacement focal spot (FWHM 22–23 mm) than the softer phantom (14–16 mm). The ARFI time constant was measured twice in each phantom. The softer Phantom 25 yielded τ = 4.0 and 4.5 ms (R” = 0.98 and 0.99), while the stiffer Phantom 50 yielded τ = 3.0 and 3.5 ms (R” = 0.62 and 0.74). In both cases the stiffer phantom exhibited a shorter time constant (Fig. 5a), and the human brain (3.70 ± 0.27 ms) fell between the two phantoms. The ARFI τ tracked the MRE-derived Kelvin-Voigt time constant: both decreased with stiffness by a nearly identical fractional amount (ARFI ratio 0.76; MRE-KV ratio 0.74; Fig. 5b), indicating that the ARFI provides a temporal mechanical contrast that was associated with independently measured viscoelastic properties in these phantoms.

**Figure 5.**
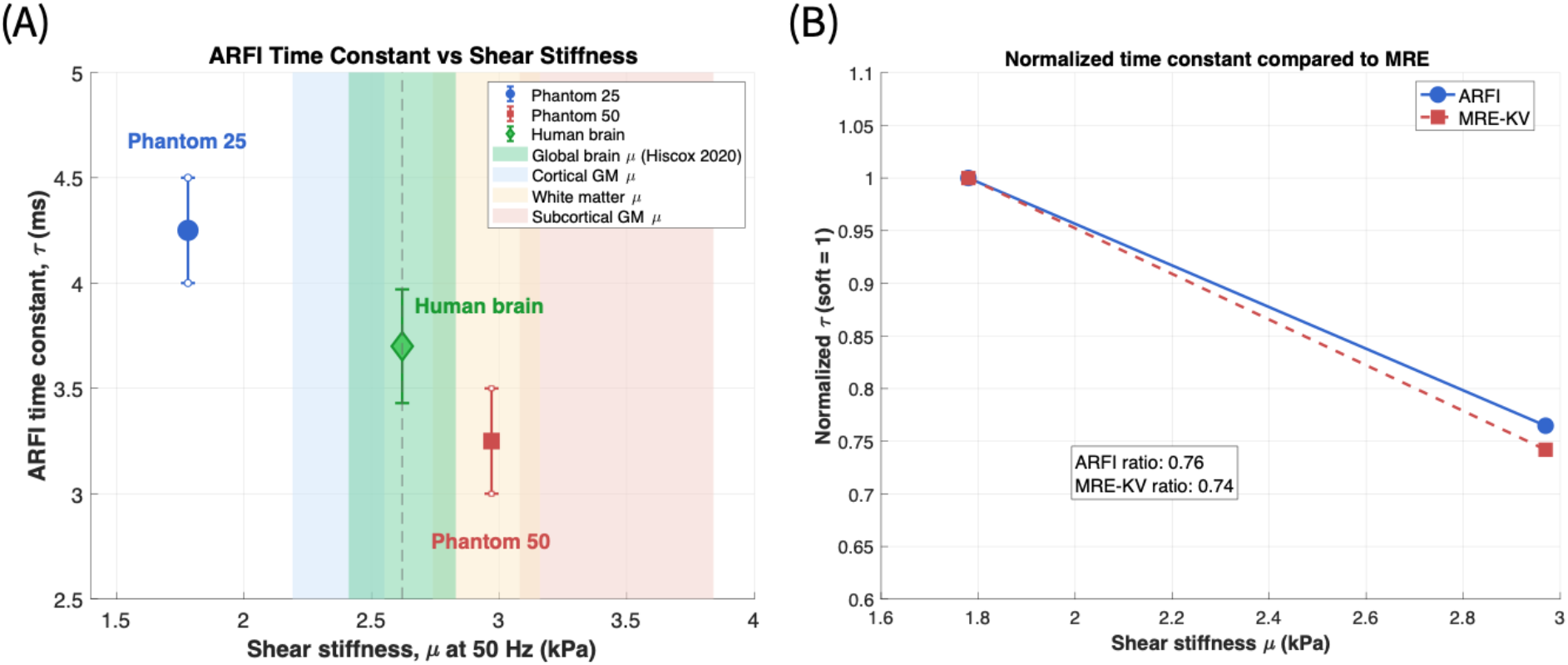
(A) ARFI τ versus MRE shear stiffness (50 Hz). Phantom points show the mean of two ARFI measurements (large symbols) with individual measurements as small open symbols (Phantom 25: τ = 4.0, 4.5 ms; Phantom 50: τ= 3.0, 3.5 ms); vertical error bars span the two measurements. The human brain (green diamond) plots the ARFI time constant measured in this study (τ = 3.70 ± 0.27 ms, mean ± SD across five participants) at the global whole-brain shear stiffness reported by Hiscox et al. (μ = 2.62 ± 0.21 kPa). Shaded vertical bands indicate the Hiscox et al. (2020) shear stiffness ranges (mean ± SD, 50 Hz) for global whole-brain (green), cortical gray matter (blue), white matter (tan), and subcortical gray matter (red); the dashed vertical line marks the global mean. (B) Normalized ARFI τ and MRE Kelvin-Voigt τ versus stiffness; both decrease with stiffness by a nearly identical fractional amount (ARFI ratio 0.76; MRE-KV ratio 0.74).

### Ultrasound Pulse Design

Simulations using the gamma-variate model illustrate how τ informs sonication pulse design (Fig. 6). For a single rectangular FUS pulse, the peak displacement increases with pulse duration relative to τ (Eq. 8), reaching approximately 80% of its maximum at T/ τ ≈ 3 and 96% at T/ τ ≈ 5 (Fig. 6, left). Given the assumed gamma-variate response and (τ = 4 ms), the model predicts that a pulse duration of approximately 12 ms to reach 80% of the maximum displacement. For pulse trains at fixed duty cycle, the pulse repetition frequency (PRF) shapes the temporal displacement profile relative to τ (Fig. 6, right): at low PRF (50 Hz), the tissue relaxes substantially between pulses, producing an oscillating displacement, whereas at high PRF (500 Hz), the displacement accumulates into a smoother, sustained response.

**Figure 6.**
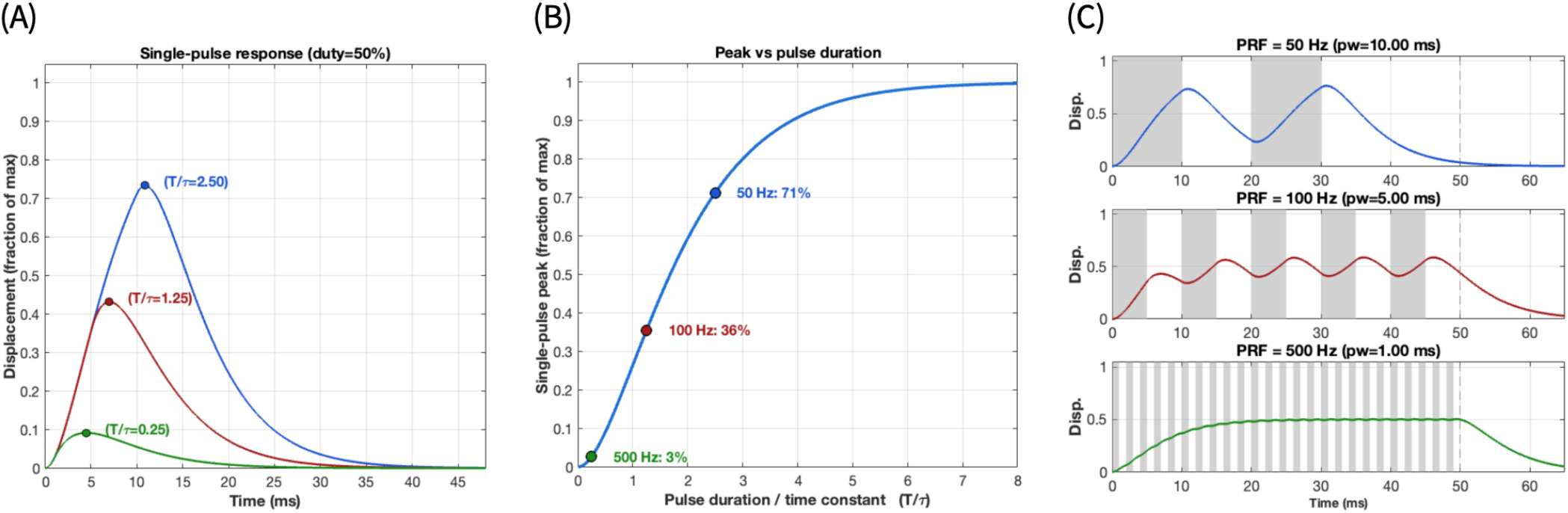
Model-derived illustration of how the time constant τ informs ultrasound pulse design (brain τ = 4 ms; duty cycle = 50%). (A) Simulated single-pulse displacement responses for the individual pulse widths corresponding to each pulse repetition frequency (PRF) at the fixed 50% duty cycle (50 Hz, T/ τ = 2.5; 100 Hz, T/ τ = 1.25; 500 Hz, T/ τ = 0.25). Longer pulses (lower PRF) produce a larger single-pulse peak (marked). (B) Peak single-pulse displacement, as a fraction of the long-pulse maximum, versus pulse duration relative to τ (Eq. 8). The three markers indicate the single-pulse peaks for the three PRF pulse widths shown in (A); displacement rises steeply for T/ τ < 2 and plateaus thereafter, reaching ∼80% at T/ τ ≈ 3, longer pulses yield little additional displacement at increasing acoustic dose. (C) Displacement time courses for pulse trains at PRF = 50, 100, and 500 Hz at the same 50% duty cycle (grey regions indicate the FUS pulses; dashed line marks the end of the train). Because the duty cycle is fixed, the time-averaged force, and hence the mean displacement level, is matched across PRFs; the PRF then governs the temporal ripple: low PRF produces large oscillations as the tissue relaxes between pulses, while high PRF produces a smooth, sustained accumulation near the mean. Thus the duty cycle sets the mean displacement while the PRF, relative to τ, determines the smoothness of the response. All curves derive from the gamma-variate model (Eqs. 1–2)

## Discussion

A prior study measured the ARF-induced displacement sonication dynamics in post-mortem pig brain using two unipolar MEGs with 10 ms FUS pulses while varying the relative timing of FUS and MEGs, fitting an exponential model to the displacement[10]. Here, for the first time, we measured the time constant τ in the living human brain. We used a bipolar MEG[18] that provides a larger dynamic range of phase shifts in both the model and the measurements, enhancing sensitivity for determining time constant, and modeled the ARF response with a gamma-variate impulse response that captures both the rise and relaxation of the displacement (Eqs. 1–2), improving upon the single exponential, which unrealistically assumes an infinitely rapid rise time of the tissue response.

Across five participants, the measurements were highly consistent (Table 1). We incorporated a time-series analysis pipeline with physiological noise regression and bulk motion correction, critical for isolating the microscopic ARF-induced phase shifts from larger motion-related noise, and a general linear model (GLM) that produces ARFI maps with statistical confidence[5] (T-scores).

To explore whether τ is sensitive to tissue stiffness, we measured τ in two custom gelatin phantoms characterized independently by MRE. The stiffer phantom exhibited a lower peak displacement (0.17 vs 0.74 μm), a broader displacement focal spot (FWHM 22 vs 15 mm), and a shorter τ (3.0–3.5 ms) than the softer phantom (4.0–4.5 ms). These observations are mutually consistent signatures of higher shear modulus, a stiffer medium deforms less (lower peak displacement) and equilibrates faster (shorter time constant). The last of these is expected from viscoelastic theory: for a Kelvin-Voigt solid, the relaxation time τ_kv_= η/μ decreases with increasing stiffness μ when the viscosity η changes less than the stiffness. Consistent with this interpretation, the ARFI τ tracked the MRE-derived Kelvin-Voigt time constant (Fig. 5b). Whereas static displacement magnitude reflects a combination of stiffness and delivered acoustic intensity, the τ is derived from the temporal shape of the response and is therefore less confounded by the intensity actually delivered to the focus, a notable advantage in transcranial applications, where in-situ intensity is difficult to determine due to skull-induced attenuation and aberration. While based on only two phantoms, this suggests that τ may provide a contrast sensitive to tissue stiffness.

Beyond tissue characterization, knowledge of the time constant has practical value for sonication pulse design. Figure 6 shows that the peak displacement produced by a single pulse depends on the pulse duration relative to time constant, while for pulse trains the interplay between PRF and τ determines whether the displacement oscillates or accumulates. Because τ characterizes how rapidly the tissue deforms and relaxes, measuring it enables sonication parameters to be tailored to the tissue’s mechanical response. For example, selecting a pulse duration that efficiently achieves a target displacement, or a PRF that produces a desired temporal profile, linking the temporal characterization presented here to the optimization of both MR-ARFI acquisition and therapeutic sonication protocols.

This study has several limitations. The sample size was small (n = 5), limiting assessment of inter-subject variability, although the consistency and high correlation across participants, despite the use of two ultrasound systems and two MRI platforms, support the robustness of the approach. Protocol modifications during data collection resulted in slightly different MEG and FUS timing across participants as experience was accumulated. The stiffness-sensitivity analysis was based on only two gelatin phantoms, and the brain stiffness used for comparison was a literature value rather than one measured in the participants. The apparent broadening of the displacement focal spot in the stiffer phantom, while reproducible, was accompanied by lower SNR, and disentangling a genuine mechanical broadening from SNR effects will require matched-SNR measurements. Regional variation in stiffness between gray and white matter is well established[12], [13], [14], [21] but tissue differentiation was not the objective of this work; extending the approach across a wider stiffness range and to regional biomechanical differences in healthy or pathological brain is an important future direction. Future studies should also examine how tissue composition, age, sex, and ultrasound parameters such as frequency influence ARF dynamics.

## Conclusion

We introduced an MR method to noninvasively and transcranially measure the human brain’s temporal dynamics in response to acoustic radiation force. This time constant τ is an essential parameter that can be incorporated into transcranial ultrasound protocols for optimizing pulse design, understanding tissue mechanical and neurocognitive properties, and tailoring therapeutic sonication toward tissue-specific, individualized treatment.

## Acknowledgments

This work was supported by NIH MH131684 and NIH EB032743.

## Conflicts of interest

No potential conflict of interest was reported by the authors.

## Data Availability

Data will be made available upon request.

## Notes

### Competing Interest Statement

The authors have declared no competing interest.

